# Linking differences in personality to demography in the wandering albatross

**DOI:** 10.1101/2025.02.27.639933

**Authors:** Joanie Van de Walle, Silke van Daalen, Samantha C. Patrick, Christophe Barbraud, Karine Delord, Henri Weimerkirch, Jack Thorley, Stephanie Jenouvrier

## Abstract

Population dynamics are shaped by individual differences. With a good understanding of the relationships between individual differences and vital rates, population models can be improved to yield more realistic and detailed demographic projections. Personality, i.e., consistent individual differences in behaviour, is expected to shape individual differences in performance. Yet, an empirical quantification of its impact on population dynamics is currently lacking. Here, we developed and analysed a three-dimensional hyperstate population model that accounts for three sources of individual differences simultaneously in its structure: age, breeding state and boldness as a measure of personality. We parameterized our model using empirical demographic and boldness data on the wandering albatross (*Diomedea exulans*) population from the Crozet archipelago. We conducted sensitivity analyses to quantify the relative importance of boldness. We then simulated scenarios of increased strength of relationship between boldness and three vital rates (survival, breeding probability and breeding success) to explore conditions under which shifts in boldness distribution could be observed in the future. We showed that the sensitivity of population growth rate to changes in vital rates followed the normal distribution of boldness within the population. Indeed, population growth rate was much less sensitive to changes in the vital rates of extreme shy or bold individuals, compared to that of individuals of intermediate boldness. Overall, sensitivity of population growth rate was greater for survival than reproductive rates across all three dimensions. Our simulations showed that increasing the strength of the relationship between boldness and survival would yield the greatest shift in boldness distribution over time compared to breeding probability and breeding success. However, shifts in boldness distribution appeared constrained by the low heritability (*<* 0.2) value and the large variance in boldness in this population. Our study provides an important contribution to our understanding of the role of personality in shaping the population dynamics of wild species. In the face of global change, our approach offers a promising avenue to predict the potential for behavioral adaptation. More generally, our approach may help to unravel the complex interplay between individual variations in any (or many) traits and population dynamics.

## Introduction

Individuals within a population do not contribute equally to population growth rate. Identifying and quantifying the importance of sources of individual variation is challenging, but crucial to adequately understand and predict population dynamics. Structured population models directly account for individual variation in their construction. Ever since the publication of Leslie (1945)’s work, the predominant structuring element in structured projection models has remained age. This is because there has been a longstanding recognition that mortality and fertility patterns often vary as individuals progress in age. However, in many systems, e.g. in invertebrates, plants and fishes, development stage often better captures the differences between individuals than age, justifying the use of stage-structured population models (Lefkovitch 1965; Caswell 2001). Morphological traits, e.g. size, also constitute a major structuring element in plant populations and have also been increasingly incorporated into animal population models (reviewed in Doak et al. 2021). For instance, size is central to integral projection models (Easterling et al. 2000). More generally, individuals can be classified by any attribute, to build models tailored to our understanding of what best accounts for individual differences in a given population. For example, the consideration of breeding states in seabirds is essential, as survival and breeding probabilities differ among breeders and non-breeders (Jenouvrier et al. 2008).

Personality, defined as consistent and heritable individual differences in behaviour (Andrew Sih et al. 2004), is ubiquitous in wild animal populations. Considerable effort has been directed towards understanding individual differences in single or multiple behavioural traits and their ecological (A. Sih et al. 2012; Smith and Blumstein 2008), evolutionary (Wolf and Weissing 2012) and conservation implications (MacKinlay and Shaw 2023). From a demographic perspective, several investigations have explored the mechanisms by which behavioral diversity is maintained within populations (A. Sih et al. 2012; Wolf and Weissing 2012) and its potential for population regulation (López-Sepulcre and Kokko 2005; Sæther and Engen 2019). This diversity can give rise to cyclic dynamics (e.g. Mougeot et al. 2003) and arise due to frequency-dependent selection, wherein one behavioral type tends to replace another in response to shifting environmental conditions. A notable example is provided by the “rock-paper-scissors” dynamics in side-blotched lizards (*Uta standburiana*), where the abundance of three morphs differing in morphology and aggressiveness levels follows cyclic dynamics in response to frequency-dependent selection (Sinervo and Lively 1996). Moreover, numerous studies have shown how density-dependent aggressiveness levels and territoriality can affect population abundance (López-Sepulcre and Kokko 2005; Bunnefeld et al. 2011; Mougeot et al. 2003).

Theoretical and empirical work suggests that demographers should be concerned with personality. First, detection probabilities of individuals expressing different personalities can vary (e.g. the increased trappability of bolder individuals; Biro and Dingemanse 2008), potentially introducing sampling bias (Biro and Dingemanse 2008). Second, there is accumulating evidence that personality is linked to migratory propensity and dispersal decisions (e.g. Clobert et al. 2009; Chapman et al. 2011; Beukeboom et al. 2023). Third, and more importantly, it has been repeatedly shown that individuals expressing different personalities can exhibit different survival and reproductive patterns (Biro and Stamps 2008; Andrew Sih et al. 2004; A. Sih et al. 2012; Smith and Blumstein 2008; Moiron et al. 2020), thereby yielding consequences for individual lifetime performances and potentially population dynamics.

Surprisingly, the influence of behavioral traits or personality on population dynamics remains under-explored empirically. This may be explained by the fact that incorporating behavioral stages, like personality, into population models poses a particular challenge because of the requirement for detailed longitudinal long-term data, and because the connections between personality and all demographic rates throughout the life cycle are rarely understood in most systems (Van de Walle et al. 2024). Yet, accounting for behavioural ecology in the construction of population models is expected to improve the accuracy of population size projections (Sæther and Engen 2019). Further, in a theoretical study, Kendall et al. (2018) demonstrated that personality-mediated differences in survival and reproduction can alter the equilibrium density of the population. Because behavioral adaptation has the potential to mitigate population declines and play a crucial role in shaping population responses to anthropogenic change (Maspons et al. 2019; Buchholz et al. 2019; Bro-Jørgensen et al. 2019), it thus appears imperative to understand and empirically quantify the demographic impact of personality within natural populations.

As long-term data accumulate and our understanding of what shapes individual differences deepens, we are now facing the challenge of having to incorporate multiple sources of individual differences (e.g. age, state, size, etc.) into matrix population models. One common practice is to aggregate information into combined classes, for example having classes defined by a combination of age classes and stages (e.g., Jenouvrier et al. 2018; Van de Walle et al. 2021). Another approach is to use multistate models, i.e., models in which individuals are simultaneously classified by two attributes. Such models were introduced by Rogers (1967) and their construction and analysis were later expanded by Hunter and Caswell (2005) through the use of the vec-permutation technique. Multistate models have been used in many systems for diverse combinations of individual attributes (e.g. Ozgul et al. 2009; Caswell and Salguero-Gómez 2013; Metcalf et al. 2012). Megamatrices are another alternative matrix construction that uses the vec-permutation approach to analyse two-dimensional models (Pascarella and Horvitz 1998).

A substantial development in matrix population models came from Roth and Caswell (2016)’s work which extended the vec-permutation approach to models with virtually any number of attributes (hereafter referred to as “dimensions”). They called such models “hyperstate matrix models”. This recent development now makes possible the integration of multiple known, or expected, sources of individual differences into a single, comprehensive, matrix population model. When projected in time, those models allow the simultaneous tracking of individual distribution along each dimension, such as age structure, distribution in the landscape, and phenotypic distribution, which can be of great interest in ecological and evolutionary studies. In addition, this model formulation facilitates the implementation of classical analytical sensitivity analyses using matrix algebra. Sensitivity analyses allow the assessment of the impact of a perturbation in each vital rate on population properties, such as population growth rate (Caswell 2019), and answer questions such as: what would happen to population growth rate if survival within or across dimensions was to change? Yet, hyperstate models have remained largely underused in natural populations. Obstacles to their general use do not lie in running and analyzing such models, but in parametrizing them because they require detailed empirical estimates of vital rates across all dimensions (Roth and Caswell 2016).

Here, we quantitatively assessed the role of personality in shaping the population dynamics of a long-lived seabird, the wandering albatross (*Diomedea exulans*) from Crozet Archipelago. Personality is heritable in this population (Patrick et al. 2013). To account for the transmission of personality traits between parents and offspring, it is essential to incorporate three interactive dimensions—personality, age, and breeding state—into the matrix population model. A hyperstate model allows us to achieve this by capturing these interactions and their influence on population dynamics. A previous study showed that the effects of boldness on vital rates were of low magnitude in this population but taken together over the entire life cycle could nevertheless affect population growth rate, especially in males (Van de Walle et al. 2024). Our main objective was to use those parameter estimates to assess the role of boldness on overall population growth rate by conducting sensitivity analyses and scenario based sensitivity analysis.

With rapid global change, individuals will be faced with novel environments. The vulnerability or success of individuals when faced with such conditions may depend on their behavioral type (Buchholz et al. 2019). For instance, individuals expressing different personality trait values may be deferentially susceptible to mortality factors, such as predation or exploitation. Sensitivity analysis can help understand how population growth rate would change if the survival, or breeding performance of individuals of a given personality type was perturbed. Thus, our first two specific objectives were to evaluate how sensitive population growth rate is to individual differences in boldness and assess through which pathway (e.g. survival) and through which component (e.g. bolder individuals) population growth rate is more sensitive. For example, with this objective we aim to answer questions such as “What would happen to population growth rate if bolder individuals had reduced survival rate, due to for example, a higher vulnerability to fishery bycatch?”. For our third objective, we aimed at verifying whether population growth rate is sensitive to the shape of boldness distribution at the population level. Finally, our fourth objective was to explore conditions under which shifts in boldness distribution could be observed and impact population growth rate.

Our main objective was to use these parameter estimates to assess how changes in boldness influence overall population growth rate and structure through a series of sensitivity analyses. With rapid global change, individuals will be faced with novel environments, and their vulnerability or success may depend on their behavioral type *\*citep{Buchholz2019}. For example, individuals expressing different personality trait values might be differentially susceptible to mortality factors, such as predation or exploitation. Sensitivity analysis can help predict how population growth rate would change if the survival or breeding performance of individuals with specific personality traits were perturbed.

To address this, we conducted three types of sensitivity analyses:

1. **Local Sensitivity Analysis**: We performed a classic sensitivity analysis to evaluate how small changes in vital rates, according to boldness, impact population growth rate and through which pathway (e.g., survival) and component (e.g., bolder individuals) these effects are most pronounced. This approach answers questions such as: “What would happen to population growth rate if bolder individuals experienced a reduced survival rate due to higher vulnerability to fishery bycatch?”
2. **Sensitivity to the Shape of Boldness Distribution**: We examined whether population growth rate is influenced by changes in the overall shape of the boldness distribution at the population level.
3. **Scenario-Based Global Sensitivity Analysis**: We explored more substantial changes in the relationship between boldness and vital rates to assess how larger shifts in these traits could affect the population growth rate.

This comprehensive set of analyses allowed us to predict changes in population dynamics resulting from any specified alterations in vital rates based on individual differences in boldness.

## Materials and Methods

### Study population and parameters

Our study is based on the long-term (1966-2022) monitoring of wandering albatrosses breeding on Possession Island (46°24’S, 52°46’E), in the Crozet Archipelago, south-western Indian Ocean. We used vital rates (*σ*: survival, *β*: breeding probability and *γ*: breeding success probability) estimated from multi-state capture-mark-recapture models applied to juveniles and adults. Personality tests were conducted on breeding adults from 2008 to 2020, which allowed the estimation of the impact of boldness on vital rates in adults (i.e., individuals that have bred at least once). Detailed methodology for the estimation of vital rates can be found in Van de Walle et al. (2024) and in Appendix S1. Also, see Appendix S2 and Patrick et al. (2013) for full description of how boldness was measured on wandering albatrosses at Crozet.

### Model description

In the wandering albatross, the vital rates vary according to age and breeding state (Van de Walle et al. 2024) and our population model included these two sources of individual variation. In addition, we included a third dimension, i.e., personality, to evaluate its importance as a structuring characteristic. To account for these three sources of individual variation simultaneously, we built an hyperstate matrix population model following Roth and Caswell (2016) to characterize individuals along three distinct dimensions, or stages: age class (*i*), breeding state (*j*) and boldness class (*k*);

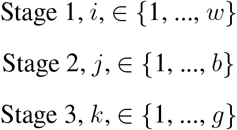

We considered transitions on an annual basis, from year *t* to year *t* + 1. We used 31 age classes (*w* = 31), with age class 1 corresponding to individuals of age 0 (fledglings). All other age classes corresponded to a year, with the exception of the last age class, for individuals aged 30+, which was left open-ended, following Rémi Fay et al. (2018).

We classified individuals into 6 breeding states (*b* = 6): Pre-breeders (PB), Successful breeders (SB), Failed Breeders (FB), Post-Successful Breeders (PSB), Post-Failed Breeders (PFB) and Non-Breeders (NB). The life cycle illustrating breeding state transitions and fertilities is presented in Van de Walle et al. (2024) and in Appendix S1 Figure S1. Transitions between breeding states are conditional on survival probability *σ*, breeding probability *β* and breeding success probability *γ*. The state PB includes individuals that have not yet bred. These individuals are of age class 1 and over. Wandering albatrosses can start to breed (lay an egg for the first time) at six years-old (R. Fay et al. 2016) and transition to the SB or FB states, conditional on whether they have successfully raised a chick or not. Although some wandering albatrosses can breed two years in a row (Barbraud and Weimerskirch 2012), most individuals skip breeding and take a sabbatical year in-between breeding events away from the colony (Tickell 1968). The states PSB and PFB include those individuals during their sabbatical year after having successfully raised a chick or not, respectively. Individuals can thus only reach the PSB and PFB states after having first reached the SB or FB states. After a sabbatical year, an individual skipping breeding again will transition to the NB state. Individuals within all breeding states at time *t* can produce a chick the following year at time *t* + 1, conditional on transiting to the SB state.

We classified individuals into 6 boldness classes (*g* = 6) of equal width, ranging from −3 to 3, with higher values corresponding to bolder classes. We chose to model six boldness classes to have a matching number of classes across dimensions (here *b* = *g* = 6). This allowed us to directly compare the relative importance of breeding state with the boldness structure for population growth rate. We could not test the scenario of an equal number of classes for boldness and age stages (i.e., *g* = *w* = 31) because of computational limitations. A model with this many classes would run, but the analytical calculation of sensitivities becomes too cumbersome.

### Notation

Here, we will briefly introduce the mathematical symbols used throughout. Matrices are denoted by upper-case boldface letters (e.g., **P**), and vectors by lower-case boldface letters (e.g., ***ρ***). Vectors are column vectors by default; **x**^T^ is the transpose of **x**. The vector **1**_*n*_ is an *n* × 1 vector of ones and **I**_*n*_ is the identity matrix of order *n*. ***e***_*i*_ is a unit vector with a 1 in the *i*^*th*^ entry and zeros elsewhere, and ***E***_*ij*_ is a matrix with a 1 in the *ij*^*th*^ entry and zeros elsewhere. The diagonal matrix with the vector ***x*** on the diagonal and zeros elsewhere is denoted 𝒟 (***x***). The symbol ° denotes the Hadamard, or element-by-element, product and ⊗ denotes the Kronecker product. The vec operator, vec **X**, stacks the columns of an *m* × *n* matrix **X** into an *mn* × 1 column vector. Block-diagonal matrices are matrices with transition matrices for a given dimension on the diagonal and zeros elsewhere are denoted by blackboard font (e.g., 𝔹). In the hyperstate model formulation, vectors and matrices have a block structure that reflects the arrangement of age classes, breeding states, and personality groups. The symbol tilde (~) is used to indicate such vectors (e.g., **ñ**) and matrices (e.g., **Ũ**). When helpful, we will indicate the dimension of matrices and vectors when they are referenced.

### Matrix construction

The population vector **ñ** has age classes organized within breeding states within boldness classes, with the entry **ñ**_*ijk*_ corresponding to the number of individuals within age class *i*, breeding state *j* and boldness class *k*;

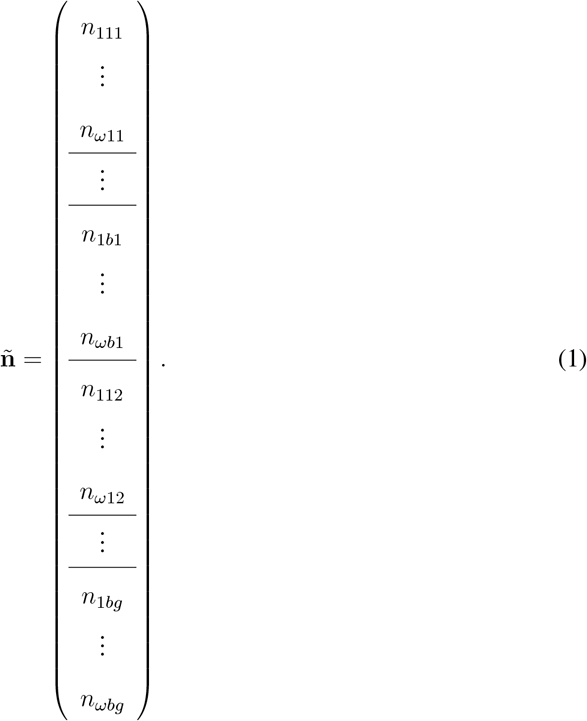

The matrix **Ã** projects the population vector **ñ** forward in time, from time *t* to time *t* + 1 and has the same structure,

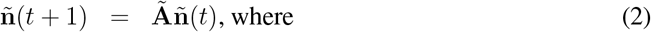

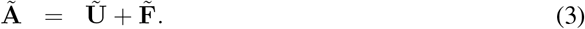

**Ã**, **Ũ**, and 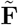 are all of dimension (*ωbg* × *ωbg*). Similar to matrix models without additional dimensions, **Ũ** contains transition probabilities for living individuals, and 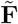 contains the production of new individuals by mature individuals. However, in hyperstate matrix formulation, transitions within **Ũ** and 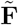 are further decomposed into distinct processes, each occurring in the distinct model dimensions (here three dimensions: age, breeding state and boldness class). For instance, **Ũ** combines three distinct processes: 1) transitions between age classes included in the block-diagonal matrix 𝕌, 2) transition between breeding states included in the block-diagonal matrix 𝕌, and 3) transitions between boldness classes included in the block-diagonal matrix ℙ. 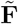 combines 1) production of offspring included in the block-diagonal matrix ℝ, 2) classification of offspring into the first breeding state included in the block-diagonal matrix 𝔽, and 3) classification of offspring into a boldness class included in the block-diagonal matrix ℍ. The decomposition of the hyperstate matrix **Ã** into its constituents is presented in Figure 1.

**Figure 1:**
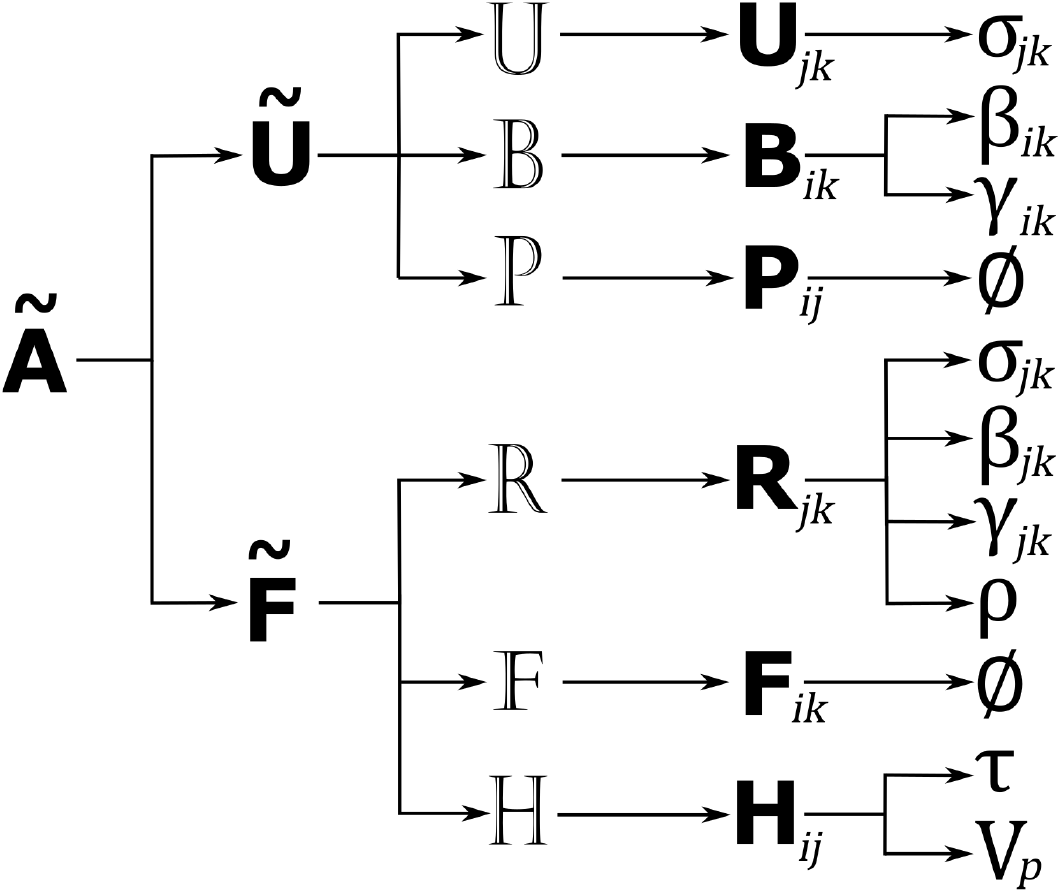
Schematic representation of the **Ã** matrix and its constituents. Note that Ø stands for empty set.

### Transitions of live individuals

The hyperstate matrix **Ũ** is obtained by sequentially multiplying the three sub-processes:

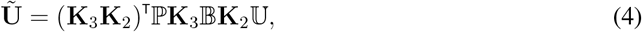

Each sub-process is captured by its corresponding block-diagonal matrix 𝕌, 𝔹, and ℙ, respectively, which are also of dimension *ωbg* × *ωbg*. Reading from right to left, individuals will move between stages at each time step following this specific sequence of transitions 1) age classes through 𝕌, 2) breeding state through 𝔹 and 3) personality class through ℙ. The vec-permutation matrices (**K**_2_, **K**_3_) rearrange the vector **ñ** for the matrix multiplication in the next dimension.

The matrix 𝕌 has matrices **U**_*jk*_ arranged on the diagonal,

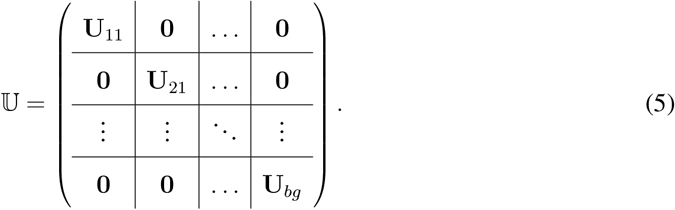

and is obtained using the following equation:

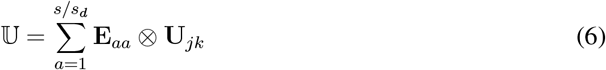

where *s* is product of dimensions, i.e., *s* = *w* × *b* × *g* and *s*_*d*_ is the size of the dimension of interest, i.e., in the case of the 𝕌 matrices, *s*_*d*_ = *w*. Note that block matrices B, P, R, F and H are constructed in the same way.

The matrices **U**_*jk*_ are of dimension *w* × *w*. For a given breeding state *j* and personality *k*, the matrix **U**_*jk*_ moves individuals between age classes *i* according to their age-, state- and personality-specific survival rates ***σ***_*ijk*_:

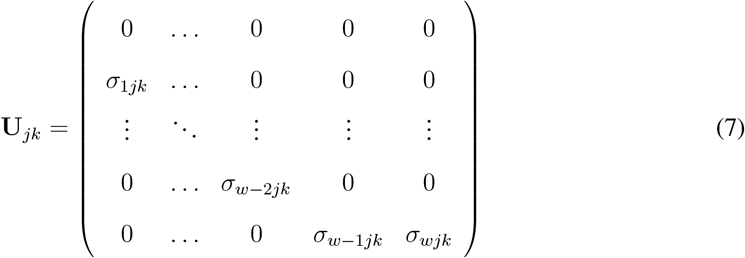

The matrices 𝔹 and ℙ have a similar structure to U as in (5), but with matrices **B**_*ik*_ and **P**_*ij*_ on the diagonal, respectively. The **B**_*ik*_ matrices are of dimension *b* × *b* and move individuals between the six breeding states (PB, SB, FB, PSB, PFB, NB) for each combination of age class (*i*) and boldness class (*k*). They have the following structure, where *β*_*ijk*_ and *γ*_*ijk*_ represent age-, state- and personality-specific breeding probabilities and breeding success probabilities, respectively:

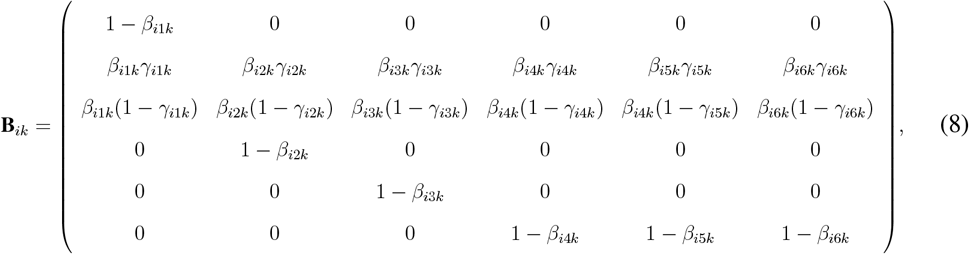

The **P**_*ij*_ matrices then move individuals from their boldness class at time *t* to a boldness class at time *t* + 1. Matrices **P**_*ij*_ are of dimension *g* × *g*. We assumed that annual transitions between boldness classes do not occur, i.e., boldness score is fixed throughout an individual’s life. This assumption is reasonable in the wandering albatross (Patrick et al. 2013). Consequently, the **P**_*ij*_ matrices are identity matrices, so that **P**_*ij*_ = **I**_*g*_ and ℙ = **I**_*wbg*_, as follows:

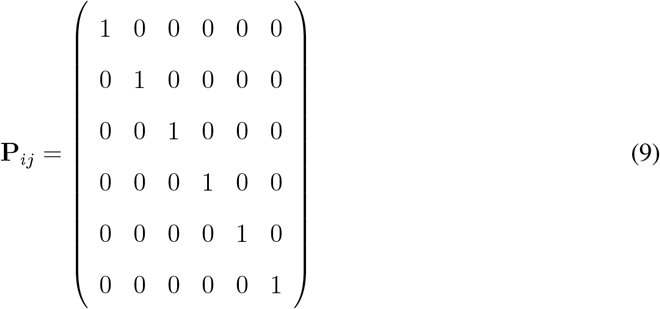

### Reproduction processes

The hyperstate matrix 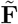 is obtained by multiplying the three sub processes:

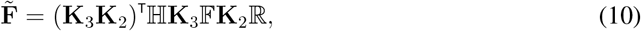

Reading from right to left, individuals will produce offspring at each time step following this specific sequence of transitions 1) production of individuals of age class 1 through ℝ, 2) classification of age class 1 individuals within the first breeding state PB through 𝔽 and 3) personality class assignation through ℍ. The vec-permutation matrices here again rearrange the dimensions of each of these matrices to respect matrix organization at each step of the multiplication.

The matrix ℝ has matrices **R**_*jk*_ arranged on the diagonal,

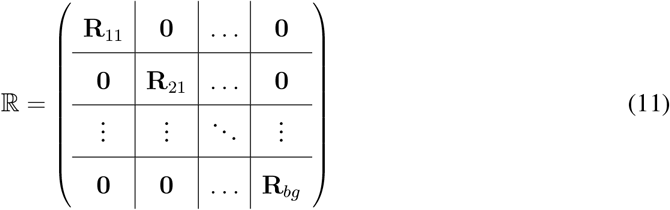

For each breeding state *j* and personality class *k*, the **R**_*ik*_ matrices are of dimensions *w* × *w* and capture the production of offspring (all in the first age class or first row) from mothers within each of the adult age classes (columns). Offspring production is dependent upon age-, state- and boldness-specific maternal survival *σ*_*ijk*_, breeding probability *β*_*ijk*_, breeding success probability *γ*_*ijk*_ and the offspring sex-ratio *ρ*, which we assumed as 0.5. The **R**_*ik*_ matrices have the following structure:

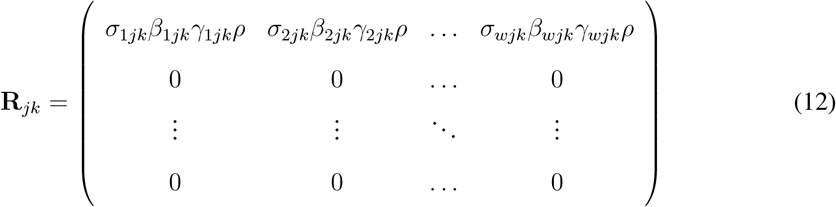

As a result of this process, all newly produced offspring are classified within the age class 1, but they remain classified within the breeding state *j* category of their mother. Reclassification of the newly produced offspring within the pre-breeder (PB) state is achieved using the matrix F. The matrix 𝔽 contains the **F**_*ik*_ matrices on its diagonal. Matrices **F**_*ik*_ are of dimension *b* × *b* and put back within the PB state the offspring produced by mothers in each breeding state :

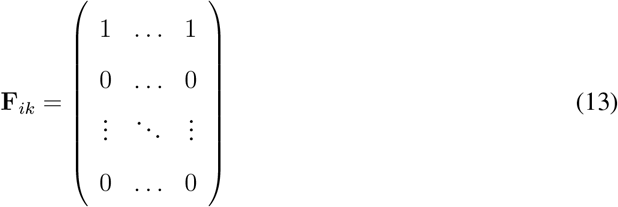

Then, the matrix ℍ classifies the newly produced offspring within a phenotypic class. The matrix ℍ contains the matrices **H**_*ij*_ on its diagonal. Matrices **H**_*ij*_ are of dimension *g* × *g* and capture the phenotypic transmission process. This process is based on a resemblance factor, *r*, corresponding to half of the heritability (*r* = *h*^2^*/*2). Heritability of boldness was estimated at 0.196 in female wandering albatrosses from Crozet (Appendix S2). Given a starting maternal phenotype, daughter phenotype was approximated using the following relationship:

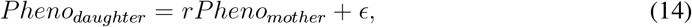

where

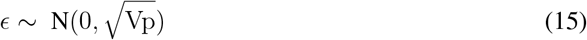

and *V p* corresponds to the observed variance in boldness score distribution at the population level, which corresponded in our case to 0.89. The distribution of observed boldness score in female wandering albatrosses is presented in Figure 5a. Each column of **H**_*ij*_ (or starting maternal phenotype *k*) consisted in a probability distribution of obtaining a given offspring phenotype (row).

## Model analysis

We calculated the asymptotic population growth rate (*λ*) as the dominant eigenvalue associated with the dominant right eigenvector, 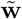 (i.e., the stable stage structure) of the population matrix **Ã** (Caswell 2001).

We analysed the model through sensitivity analyses and simulations. Boldness has a limited impact on female vital rates in the wandering albatross, which could be argued as being a limiting factor for detecting demographic impacts of boldness with a female-only model. In males, a negative impact of boldness on breeding probability was documented (Van de Walle et al. 2024). Therefore, in parallel we also analyzed a male-only model and present the results in Appendix S4.

### Sensitivity analyses

We performed sensitivity analyses to explore the relative importance of age-, state- and boldness-specific vital rates for population growth rate, *λ*. As described in Roth and Caswell (2016), in hyperstate population models, the dynamics of the population depends on the matrix **Ã**, which in turn depends on the block-diagonal matrices (e.g. A). The block-diagonal matrices depend on the dimension-specific matrices (e.g. **A**), which in turn depend on the vector of parameters of interest ***θ***. In our case such a vector contains the age-, state- and boldness-specific vital rates, and we can trace the dependency of **Ã** to such vectors through **Ũ** and 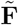 and their components (Figure 1). Following Roth and Caswell (2016) and matrix calculus rules (Magnus and Neudecker 1985), we calculated the sensitivity of *λ* with respect to, for example, ***σ***_*jk*_ as follows:

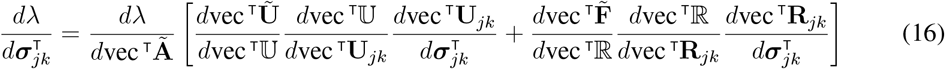

which gives:

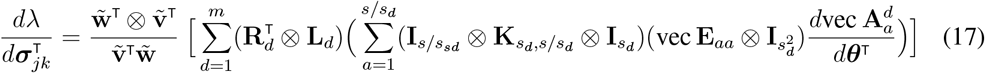

where 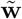 and 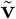 correspond to left and right eigenvectors of to the maximum eigenvalue *λ*. **R**_*d*_ and **L**_*d*_ correspond to the segment on the right- or left-end side of the block-diagonal matrix corresponding to the sub-process (or dimension) of interest in equation 4 or 10 (depending on the process in which the parameter is implied), respectively. For example, when focusing on the age dimension (d = 1), *s*_*d*_ = *ω* = 31 and the term *s/s*_*d*_ = 60. This means that within dimension 1, there are a total of 60 **A** matrices (one for each combination of state *j* and boldness score *k*). More details on the derivation of parameters within the dimension-specific matrices **A** can be found in Appendix S3. We present results in terms of elasticities, i.e., relative sensitivities, which was calculated by dividing elasticities by parameter values.

### Simulations

We simulated the impact of boldness distribution at equilibrium on population growth rate by perturbing the stable-stage distribution 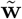 vector associated with **Ã**. We considered a scenario where the distribution of boldness at equilibrium was uniform. We then recalculated the sensitivities of *λ* with respect to the *θ* vector using the perturbed 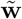.

Then, we assessed the impact of heritability on population dynamics in two ways: 1) we compared the initial distribution of boldness with that obtained at equilibrium using observed parameter values and 2) we simulated different strengths of relationship between mother and daughter phenotypes – through changes in the heritability parameter *h*^2^ - and compared the resulting boldness distribution at equilibrium. The distribution of boldness scores at equilibrium was obtained by summing the proportions of individuals within each boldness class across the other two dimensions given in the vector 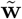. We simulated values of *h*^2^ ranging from 0 to 1 with increments of 0.25. Heritability value of 0 and 1 mean that boldness of daughters is randomly drawn within the observed distribution of boldness or is half that of their mother, the other half – i.e., male’s contribution - is randomly drawn, respectively. In addition, because we wanted to test the sensitivity of our results to our parametrization of heritability in the matrix model, as an extreme case we also considered a scenario of clonal reproduction where mothers transmit exactly their phenotype to their daughters. In all simulations, we kept the effect of environmental noise in daughter phenotype attribution.

Lastly, we simulated scenarios of different magnitudes of the functional relationship between boldness and vital rates (i.e., strength of selection) to further explore conditions under which shifts in boldness distribution could be observed and have demographic consequences. We considered scenarios where boldness would have a positive impact on vital rates in our explorations. Opposite scenarios would yield similar conclusions as to relative magnitude of boldness impacts on population dynamics. We simulated six scenarios with slopes between vital rates and boldness varying from 0 to 0.5 (logit scale) with increments of 0.1. For each scenario, we tested five heritability values ranging from 0 to 1 with increments of 0.25. For each simulation, we calculated asymptotic population growth rate (*λ*) and mean boldness value at equilibrium.

## Results

Under observed parameter values, population growth rate at equilibrium *λ* was 1.018. Across all age classes, breeding states and boldness classes, elasticity of *λ* was greatest with respect to *σ* (survival), followed by *β* (breeding probability) and *γ* (breeding success; Figure 2). Elasticities of *λ* with respect to all vital rates was high in the age class 31 (Figure 2a) because this last age class consists of individuals of age 31 and over and thus includes more individuals relative to the other single age classes. Elasticity of *λ* was also greatest for survival of pre-breeders (PB), who are the sole representative of age classes *<* 6. Perturbing survival and breeding parameters for post-successful breeders PSB would affect population growth rate more strongly than any other adult breeding state (Figure 2b). Individuals in this state are in their sabbatical year and have a great reproductive potential; their survival and return to the breeding grounds to produce a fledgling brings a disproportionate contribution to population growth.

**Figure 2:**
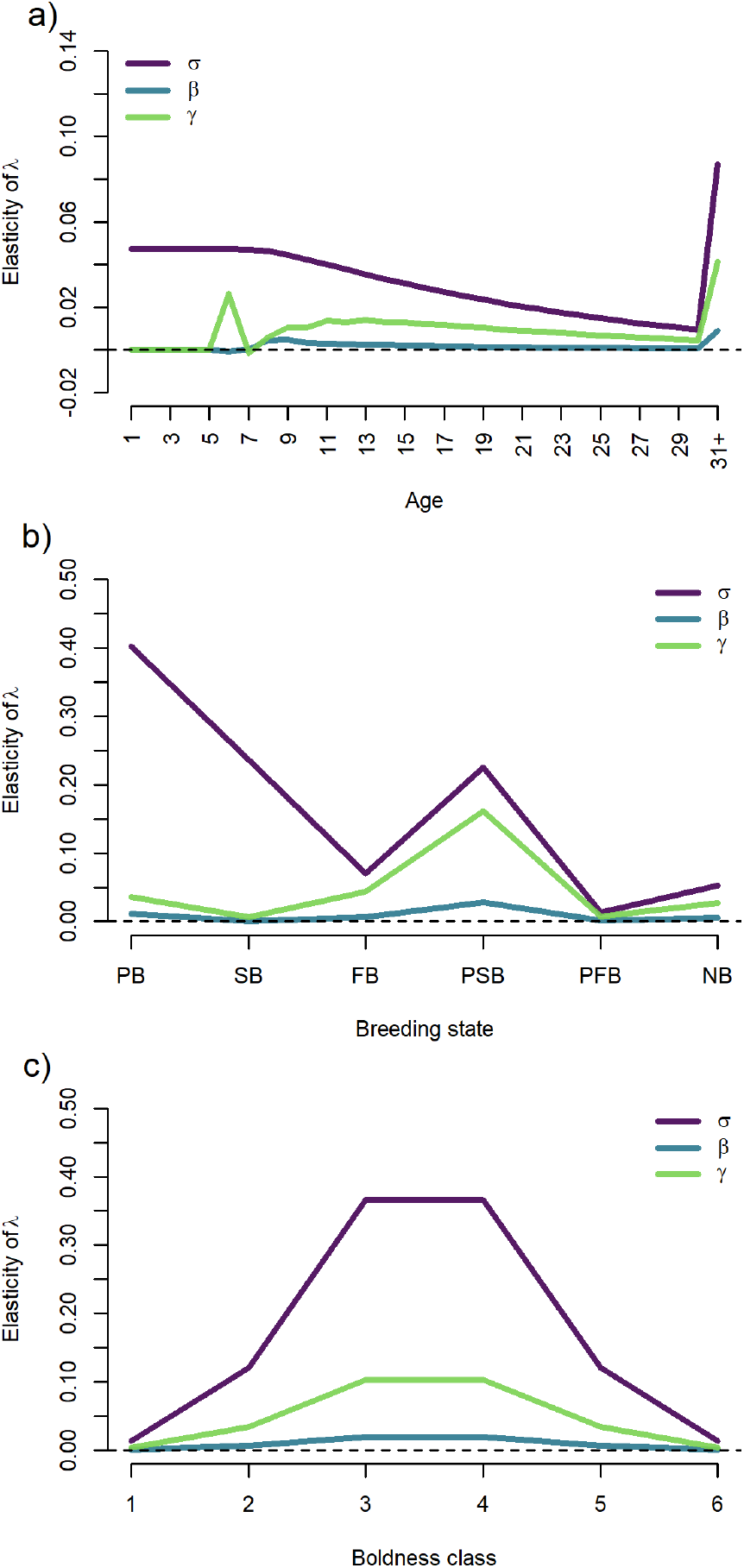
Elasticities (relative sensitivities) of population growth rate, λ to vital rates (survival *σ*, breeding probability *β* and breeding success probability *γ*) along the three dimensions included in the hyperstate model: age class, breeding state and boldness class in female wandering alabrosses at Possession Island. Note that the age class 31+ includes all individuals of age 30 years-old and over, hence the drastic increase in elasticities for this age class.

Elasticities of *λ* to all three vital rates showed a bell shape along boldness classes (Figure 2c), with higher values around intermediate boldness classes, which comprises a greater proportion of the population. This indicates that perturbing survival and breeding parameters of individuals of average boldness scores would have the greatest impact on population growth. Overall, we found that the magnitude of differences in elasticity values (Figure 2c) was similar across boldness classes to what was found across breeding states (Figure 2b), indicating that accounting for individual differences in boldness is similarly important as accounting for differences in breeding states for the population dynamics of wandering albatrosses.

When analyzing patterns of elasticities at a finer scale, we found that *λ* was most sensitive to perturbations in survival rates of pre-breeders, but only for age classes *<* 8 (Figure 3a). For age classes 8 and over, changing survival of successful breeders and post-successful breeders would yield the greatest impact on *λ*. Increasing breeding probabilities *β* of pre-breeders of age class 6 would have a negative impact on population growth (Figure 3b). This is because those individuals starting to breed at age class 6 will become either SB or FB at age class 7 and then either PSB or PFB at age class 8, where survival is very low (0.66; Table S2). Increasing breeding probabilities of PB would have the greatest positive impact on population growth in age classes 7 and 8 (Figure 3b), after this the benefit for population growth rate of increasing breeding probability for pre-breeders declines with age. Higher breeding probabilities and breeding success for PSB would also have strong impacts on population growth rate, especially at prime ages (ages 10-20; Figure 3b,c). Interestingly, despite a negative impact on population growth rate of increasing breeding probabilities of age class 6 individuals, an increase in breeding success in that age class would have a strong positive impact. This may mean that if individuals start reproducing early, they should be successful in doing so to have a positive demographic contribution. However, increasing breeding success of pre-breeders of age class 7 would have a negative impact on population growth rate because it means more individuals will reach the state successful breeder early, where breeding success is low.

**Figure 3:**
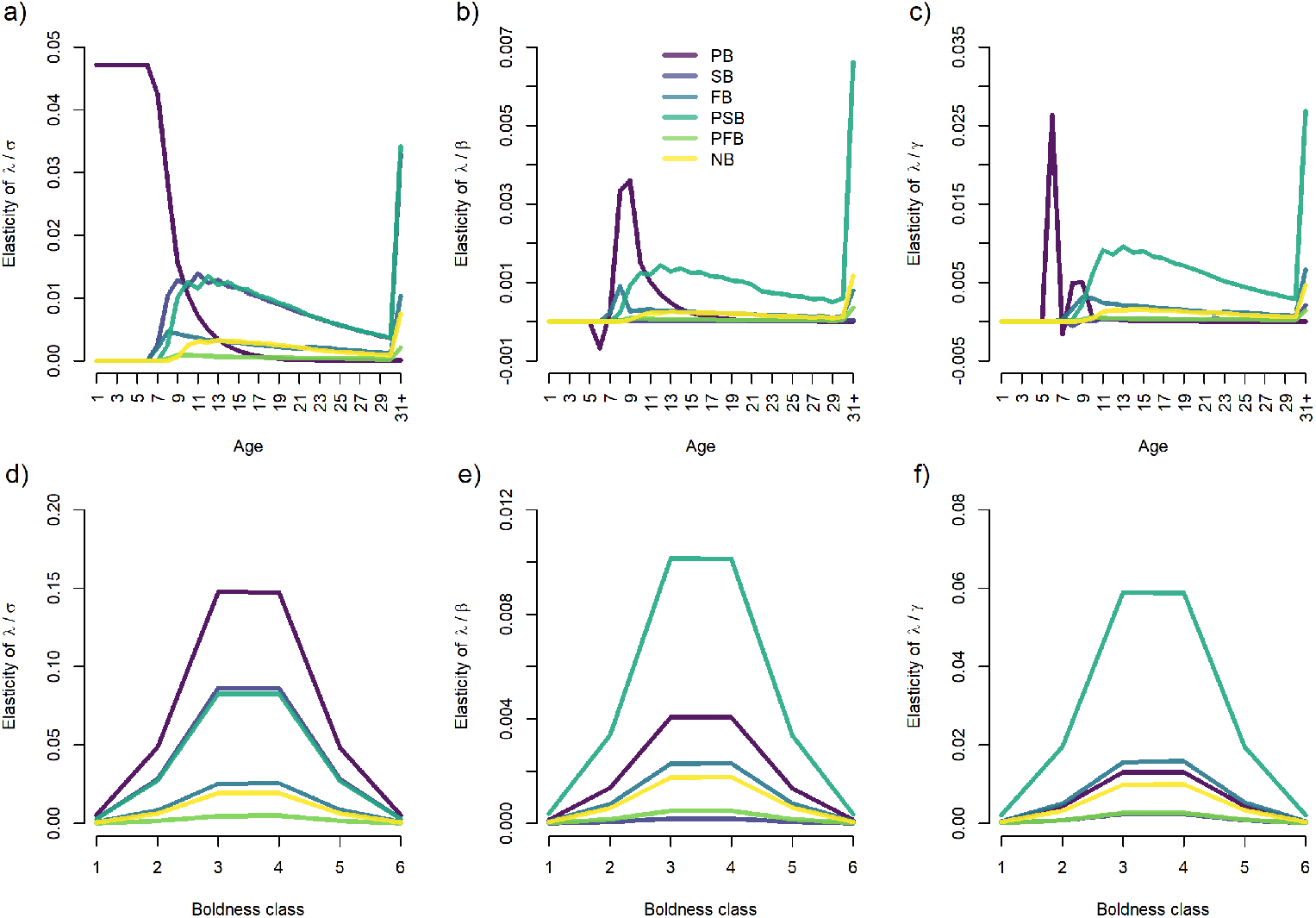
Elasticities (relative sensitivities) of population growth rate, λ to vital rates: survival *σ* (left column), breeding probability *β* (middle column) and breeding success probability *γ* (right column) for female wandering albatrosses at Possession Island.

The pattern of elasticities of *λ* to vital rates analyzed per breeding states also had a bell shape, with greater values at intermediate boldness scores (Figure 3a-c). However, results varied between survival and breeding parameters. The greatest elasticities were found for pre-breeders of intermediate boldness classes for survival (Figure 3d) and for post-successful breeders of intermediate boldness classes for breeding probability (Figure 3e) and breeding success (Figure 3f).

Assuming a uniform distribution of boldness at equilibrium and recalculating the elasticities of *λ* to vital rates, we found that overall, changing the distribution of boldness at equilibrium to a uniform distribution showed that elasticities varied very little across boldness classes (Figure 4). This indicates that the bell shape patterns of elasticities observed at Figures 2c and 3d-f result from the initial normal distribution of boldness score at the population level. Indeed, the initial distribution of boldness and that observed at equilibrium are similar (Figure 5a). Changing the value of heritability would only slightly change the distribution of boldness at equilibrium by mostly reducing the proportion of individuals within the intermediate boldness classes (Figure 5b). Only under an extreme scenario assuming clonal reproduction would we observe a marked change in the boldness distribution towards a increased proportion of shyer and bolder individuals.

**Figure 4:**
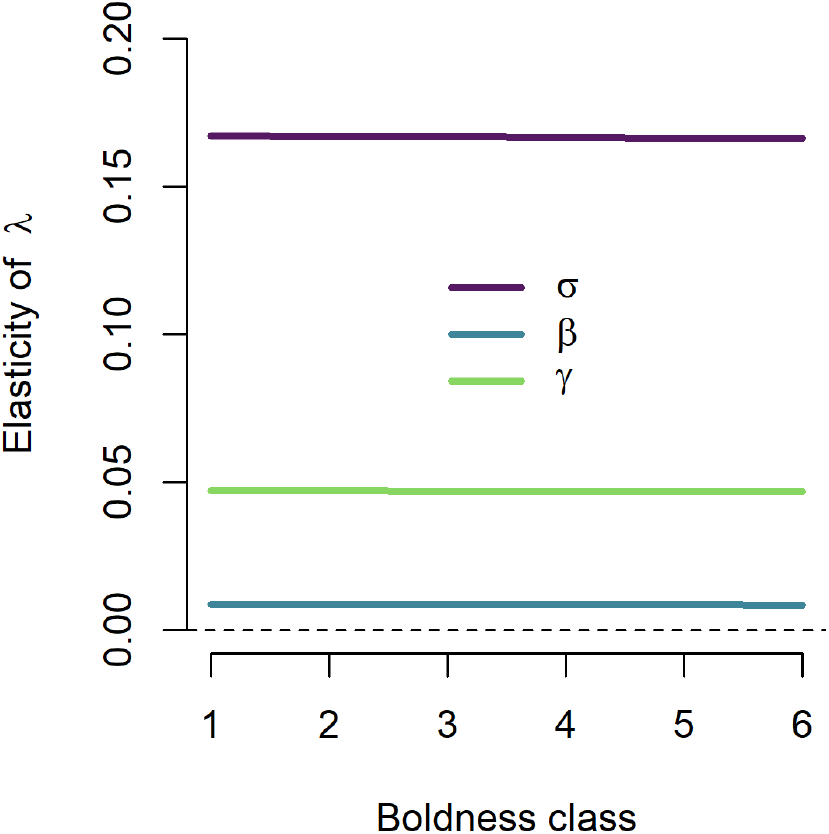
Elasticities (relative sensitivities) of population growth rate, *λ* to vital rates: survival *σ*, breeding probability *β* and breeding success probability *γ* for female wandering albatrosses at Possession Island, assuming a uniform distribution of boldness at equilibrium.

**Figure 5:**
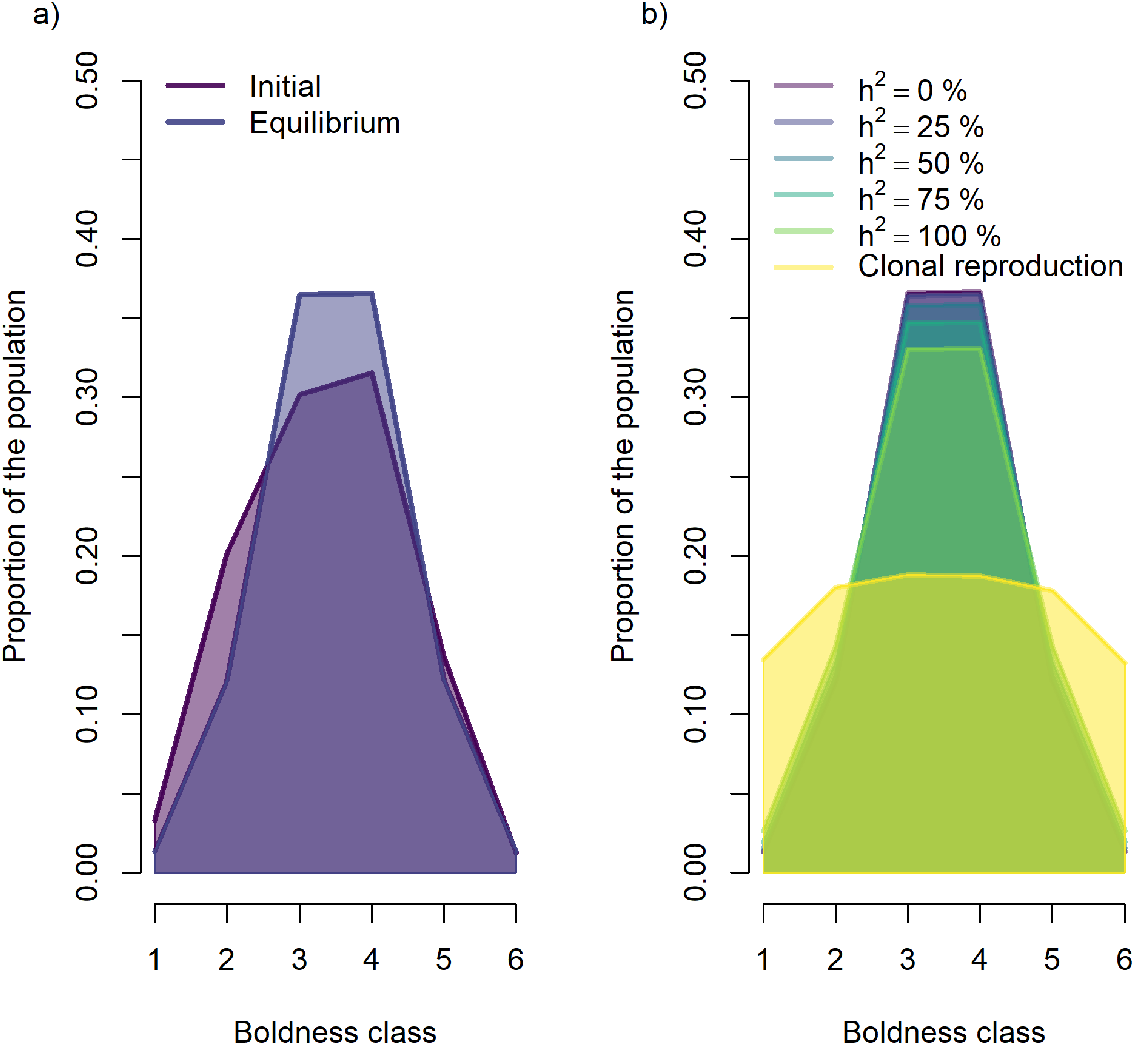
(a) Proportion of the female wandering albatross population within each boldness class observed (initial distribution) and at equilibrium using estimated parameters, and (b) proportion of the population within each boldness class using different heritability values. Under the scenario of clonal reproduction, we assumed perfect correlation between mother and daughter phenotype, but accounted for environmental variance in daughter phenotypes.

Further, for different values of heritability, here we tested six hypothetical scenarios of variations in the strength of the relationship between boldness and vital rates (Figure 6a,d,g). For each scenario, we inspected the mean boldness score at equilibrium and population growth rate. We found that varying the slope of the regression between boldness and survival would yield the greatest impact on mean boldness score and population growth rate, compared to breeding probability and breeding success (Figure 6). Increasing heritability value had a lower impact on population growth rate and mean boldness score compared to increasing the slope of the relationship between boldness and vital rates. However, population growth rate and mean boldness score increased more rapidly with increasing heritability when the slope of the relationship was higher. The ranges of difference between the minimum and maximum values for boldness score and population growth rate were small across all simulated values for survival (i.e., boldness score: 0.066; population growth: 0.004) and were negligible for breeding probabilities and breeding success.

**Figure 6:**
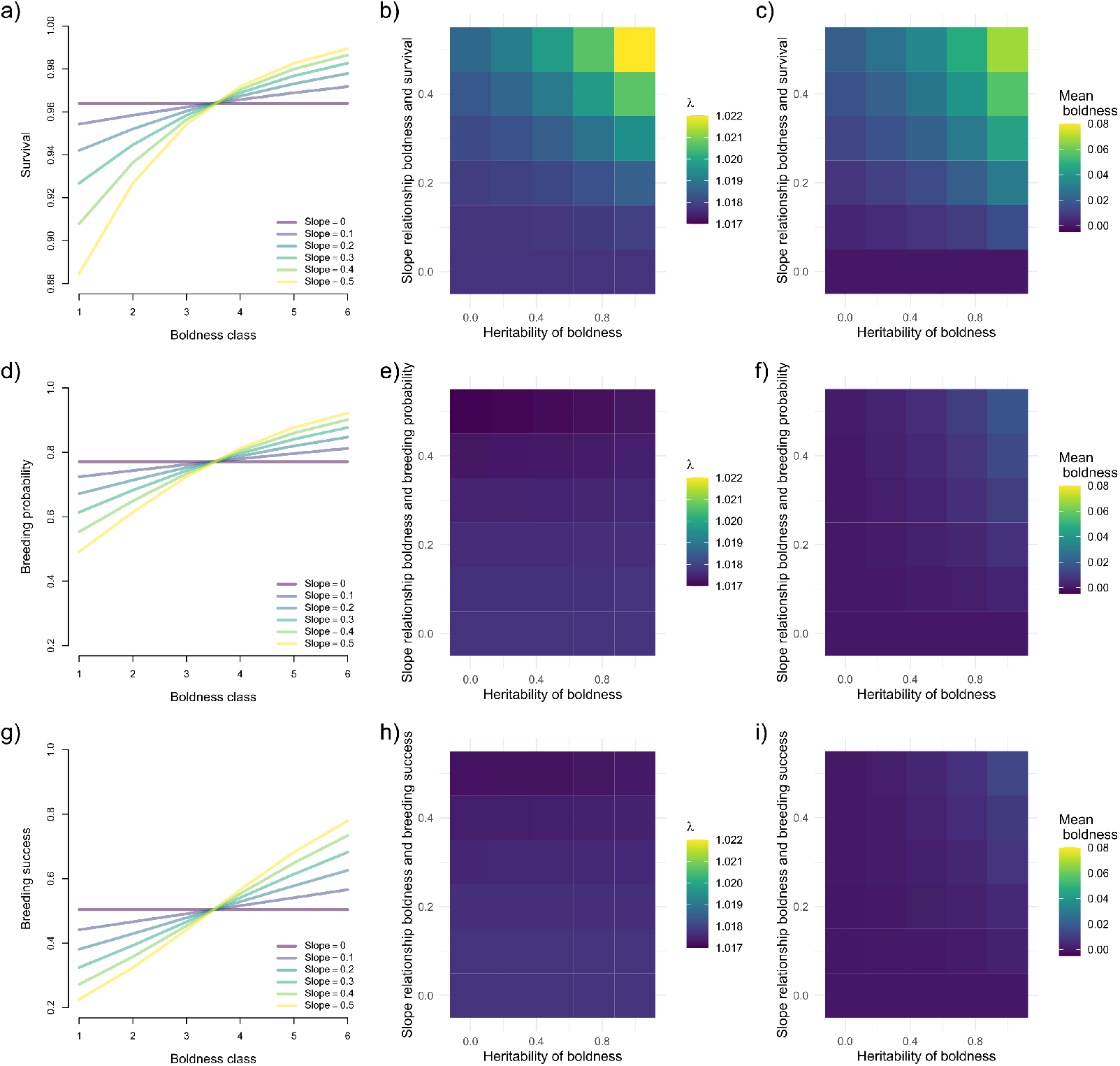
Interactive effects of heritability value and slope of the regression between boldness and survival on population growth rate in the wandering albatross at Possession Island. The left panels (a, d, and g) show the six functional relationships between boldness and the three vital rates (survival, breeding probability and breeding success) tested. The center (b, e, and h) and left (c, f, i) panels show the impact of the different slopes (y axis) and heritability values (x axis) combinations on population growth rate (λ) and mean boldness value at equilibrium, respectively. Boldness values were calculated by multiplying the number of individuals within each boldness class at equilibrium with the mid value of each boldness class.

## Discussion

Our main objective was to evaluate whether individual differences in personality can affect population dynamics. We used the hyperstate matrix formulation (Roth and Caswell 2016) to develop a three-dimensional population model structured by three attributes: age, breeding state and personality. We applied this model to empirical data from the long-term demographic monitoring of the wandering albatross from Crozet and used boldness as a measure of personality. We tested for the relative sensitivity of population growth rate to changes in vital rates of individuals across the shy-bold axis of variations and made projections of population growth rate and average boldness score under various pathways of selection. Our work provides an empirical demonstration that accounting for individual differences in personality matters from a demographic standpoint, even when their impact on single vital rates is limited. This is because perturbations of individual vital rates have more impact when directed towards the most common phenotype classes. We applied the model to boldness, but our approach would be suitable for any other personality trait, or more generally, to any phenotypic trait.

In a simulation study, Kendall et al. (2018) assumed that boldness mediated aggressiveness and a trade-off between survival and reproduction, following expectations from the pace-of-life hypothesis (Dammhahn et al. 2018; Réale et al. 2010). They then theoretically explored the impact of boldness on population dynamics using a population model with two morphs: bold (aggressive) and shy (nonaggressive) individuals. They predicted that accounting for individual differences in boldness could have an impact on morph frequencies and population abundance at equilibrium, but they advocated the necessity of using empirical data to verify their predictions. In our study, we directly incorporated individual differences in boldness into a population model parameterized using empirical data and found a limited impact of boldness-mediated differences in vital rates on equilibrium distribution. This may be explained by the absence of evidence that boldness correlates with a trade-off between survival and reproduction in the wandering albatross (Van de Walle et al. 2024), which is in accordance with limited support for the pace-of-life syndrome in empirical studies in wild animal species (Moiron et al. 2020).

Our results revealed that the equilibrium distribution of boldness plays an important role in population dynamics. Indeed, the equilibrium distribution of boldness was bell-shaped, and so was the pattern of elasticities across boldness classes. We verified this by forcing a uniform distribution of boldness at equilibrium and showing that under such a distribution, the pattern of elasticities was then mostly uniform across boldness classes. Population growth rate is thus most sensitive to changes in vital rates amongst the most frequently represented boldness class at equilibrium. This is because 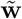, the stable stage distribution (or equilibrium distribution), is directly involved in the calculation of sensitivities. However, this equilibrium depends on the relative benefits of bolder versus shyer individuals. Indeed, when boldness confers no, or very little, fitness benefit (a scenario similar to what we observe in the wandering albatross), perturbations in vital rates of average boldness classes would lead to a greater change in population growth rate than if similar changes in vital rates were to occur amongst more extreme boldness classes. In contrast, if the fitness benefit of bolder individuals increases, through e.g. increased survival, and if the heritability of boldness is high, then boldness distribution at equilibrium would shift towards bolder individuals. Then, the bell-shaped pattern of elasticities would shift towards bolder classes, indicating that population growth rate would become more sensitive to changes in vital rates amongst bolder classes of individuals. This scenario is presented in Appendix S4 (Figs S6-S7). Alternatively, under a scenario where bolder individuals would experience reduced survival, as is typically assumed (Réale et al. 2010; Kendall et al. 2018), opposite predictions could be made. Regardless, observing shifts in boldness distribution that would impact population growth rate is unlikely in this population because it would require a combination of strong selective pressure and heritability value. Because of this, our results suggest that the wandering albatross population of Crozet should be relatively buffered against selection acting on either end of the shy-bold continuum.

Including demographic information in more than one dimension in a population model allows for quantifying sensitivities at a much finer resolution (Caswell and Salguero-Gómez 2013). Here, in addition to revealing the patterns of sensitivities across personality classes, we also found interesting results across age and breeding states worth mentioning. Indeed, our results revealed that within and across dimensions, population growth rate was very sensitive to perturbations in pre-breeder survival, especially for age classes 7 and below, after which sensitivities declined rapidly. All individuals of younger age classes are pre-breeders, and only upon age class 7 can individuals be in other breeding states. Thus, a large proportion of the population is within the pre-breeder state, which may explain the large elasticities for this breeding state. Also, juveniles can only contribute to the population if they survive to recruitment, hence reducing their survival would greatly reduce population growth rate. The combined elasticities of population growth rate to survival across all adult breeding states nevertheless outpaces those of pre-breeders (juveniles).

This result is in line with the expectation and empirical evidence showing that for long-lived species population growth rate is more sensitive to changes in survival in adults compared to juveniles (Gaillard and Yoccoz 2003).

Moreover, our analysis revealed that early reproduction might have a negative impact on population growth rate. Allowing more pre-breeders to breed for the first time at age class 6, i.e., the earliest age for first reproduction in our population (Rémi Fay et al. 2015), leads to a reduced future survival probability. This might appear contradictory to results from (R. Fay et al. 2016) showing that individuals recruiting earlier had a greater probability to become higher quality (sensu Nussey et al. (2008)) adults, i.e., have higher breeding probabilities, breeding success and survival. However, individuals recruiting as six years old had lower success compared to individuals recruiting later (R. Fay et al. 2016). Our demographic analysis shows that in fact, an increase in recruitment rate at age class six would be detrimental when considered at the population level. In wandering albatrosses, breeding is conditional on reaching a threshold body mass (Weimerskirch 2018). Even though some individuals may attain this threshold at six years old (2% of the birds; Rémi Fay et al. (2015)), this seems to come at the cost of a short-term reduction in breeding success and survival. Considering that body mass has increased in this population over the last decades (Weimerskirch et al. 2012), we can expect a shift towards earlier reproduction. Our results thus highlight the fact that such a change is likely to impact the population dynamics of the wandering albatross.

Our findings emphasize that changes in the equilibrium distribution of boldness traits become noticeable primarily under significant selective pressures. This observation becomes particularly significant when strong relationships between maternal and daughter phenotypes come into play. It’s worth noting, however, that these shifts have a limited influence on the overall population growth rate. It is possible that the limited impact of personality on population dynamics in our study stems from the little influence of boldness on female vital rates. To delve deeper into these dynamics, we conducted simulations using an extreme scenario characterized by high heritability and a much stronger link between survival and boldness. This could represent an hypothetical scenario where bolder wandering albatrosses would have better access to food resources. The inverse relationship could represent a scenario of increase vulnerability to mortality factors, such as fishery bycatch, for bolder individuals. These simulations unveiled an amplified sensitivity of population growth rate to variations in survival rates of bolder individuals.

In males, boldness was found to have a more decisive impact on vital rates; bolder males reproduced less frequently (Van de Walle et al. 2024). Overall, because they skipped reproduction more, bolder males had lower lifetime reproductive success compared to their shyer counterparts. However, our sensitivity analyses and simulations applied to males yielded very similar patterns to females (Appendix 3) and considering the lower relative sensitivity of population growth rate to reproductive rates compared to survival in long-lived species (Gaillard and Yoccoz 2003), a stronger impact of boldness on breeding probably has no noticeable impact on population growth rate overall.

Our model facilitated the examination of potential shifts in boldness distribution under varying conditions of heritability and selection strength. Our findings indicate that, based on our empirical data, the evolutionary potential of boldness is limited in the wandering albatross. This could be attributed to the species’ extended generation time, which limits rapid evolution over short time scales. Testing other species with contrasted generation times would help clarify this possibility. Also, the low heritability of boldness and large environmental variance in offspring boldness attribution may limit the evolutionary potential of boldness in this population. Additionally, this phenomenon could stem from the way we parametrized the transmission of phenotypes from mothers to daughters. We acknowledge that incorporating principles of quantitative genetics would improve model realism and predictions of evolutionary response. Introducing an additional dimension into the hyperstate model formulation could potentially enable tracking changes in both breeding values and phenotypes. However, the focus of the present study revolved around demography rather than evolution, and extending our already complex model to a four-dimensional model would pose a significant computational challenge.

In this study, we conducted a quantitative examination of the impact of boldness on the population dynamics of wandering albatrosses in Crozet. Boldness is assumed fixed over an individual life in this population, hence we included no dynamic progression in this trait in the model. Using an hyperstate model formulation in this case could be perceived as an unnecessary complication. However, using this formulation provided many advantages, such as a simplified quantification of sensitivities, and this, along each dimension separately, as well as the inclusion of a function to account for trait heritability. Importantly, the hyperstate matrix formulation holds broader applicability, extending beyond this specific species and trait. This inclusive approach opens avenues for addressing various ecological and evolutionary questions. For instance, while our study assumed a fixed phenotype (boldness) throughout an individual’s life and across varying environmental conditions, this does not need to be the case. In fact, the flexible model construction permits the incorporation of environment-dependent transitions between phenotypic classes, a valuable feature when investigating situations involving phenotypic plasticity (Childs et al. 2016). The primary prerequisite and potential constraint lies in the quantification of the relationships between phenotypic traits and vital rates over the entire life cycle of a species to construct the dimension-specific transition matrices.

In conclusion, we empirically investigated the influence of personality on the population dynamics of a wild species. While our investigation delved deeply, we observed limited effects, even when simulating more pronounced impacts of personality on vital rates. The extent to which these results can be extrapolated to other species remains an open question that requires further exploration. The intricate interplay between behaviors and demography is an active, but still underexplored research area. Consequently, more studies are needed to unravel the complexity and subtlety of behavioral effects on population dynamics, especially as populations will have to adapt to novel environmental conditions (Maspons et al. 2019).

## Notes

### Competing Interest Statement

The authors have declared no competing interest.

